# M3C: Monte Carlo reference-based consensus clustering

**DOI:** 10.1101/377002

**Authors:** Christopher R. John, David Watson, Dominic Russ, Katriona Goldmann, Michael Ehrenstein, Costantino Pitzalis, Myles Lewis, Michael Barnes

## Abstract

Genome-wide data is used to stratify patients into classes for precision medicine using clustering algorithms. A common problem in this area is selection of the number of clusters (K). The Monti consensus clustering algorithm is a widely used method which uses stability selection to estimate K. However, the method has bias towards higher values of K and yields high numbers of false positives. As a solution, we developed Monte Carlo reference-based consensus clustering (M3C), which is based on this algorithm. M3C simulates null distributions of stability scores for a range of K values thus enabling a comparison with real data to remove bias and statistically test for the presence of structure. M3C corrects the inherent bias of consensus clustering as demonstrated on simulated and real expression data from The Cancer Genome Atlas (TCGA). For testing M3C, we developed clusterlab, a new method for simulating multivariate Gaussian clusters.

## Introduction

Stratified medicine is the concept that patients may be clustered into classes to personalise patient therapy. Increasingly, patient genome-wide expression data is being used to perform clustering^1-6^. Cluster analysis of genome-wide data (e.g. transcriptomics, epigenomics, proteomics, and DNA copy number) has been shown to identify tumour subtypes with distinct clinical outcomes in cancer research^1-6^, and is starting to be applied on other diseases as well^7-9^. Therefore, there is high demand for methods that deliver robust results. Broadly, the clustering problem may be broken down into two steps: select K and separate the data into K groups. The order of these steps varies by clustering algorithm – K must be defined upfront in ***k***-means, for instance, while it is defined afterwards in hierarchical clustering. In this study, our primary focus was to develop a method for estimating the optimal K.

Numerous methods have been proposed for estimating K, such as: Monti et al. consensus clustering^10^, the GAP-statistic^11^, CLEST^12^, and progeny clustering^13^. The concept behind consensus clustering is that the ideal clusters should be stable despite resampling. Therefore, the degree of cluster stability for each value of K can be measured to estimate the optimal K. Șenbabaoğlu et al. made a useful contribution by demonstrating that false positive structures could be found in K=1 null data using the Monti consensus clustering algorithm^14^, this is a common problem in cluster analysis. The authors suggested to generate null datasets with the same gene-gene correlation structure as the real data to evaluate cluster strength. However, they did not provide a method for performing a formal hypothesis test. They developed a new metric that measures cluster stability called the proportion of ambiguous clustering (PAC) score, this is better able to estimate K than the original delta K metric^10^ proposed by Monti et al. However, the PAC score does not take into account null reference distributions, has inherant bias towards higher values of K, and does not test the null hypothesis K=1.

Our aim was to solve these problems by enhancing the Monti consensus clustering algorithm to include a Monte Carlo reference procedure to eliminate bias towards higher values of K and to test the null hypothesis K=1. This method we call M3C (https://www.bioconductor.org/packages/3.7/bioc/html/M3C.html). To introduce M3C, it is instructive to define the hypotheses that it tests. M3C calculates null distributions of PAC scores for each K (starting with K=2) by simulating K=1 null datasets. For each K, this allows us to formally test the following null hypothesis:

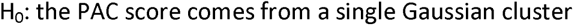

The alternative hypothesis tested for each K is:

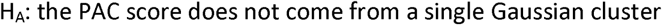

If no p values are significant along the range of K we accept the null hypothesis H_0_ in every case, this means there is no significant evidence for clusters in the data. If a p value is significant, then we can reject the null hypothesis H_0_, thereby accepting H_A_, this is significant evidence for clusters in the data. M3C presented us with an opportunity to test two hypotheses on real data. First, that pre-existing high-profile publications contain results that declare evidence of structure when in fact there is none. Second, that not considering reference distributions when deciding K leads to systematic bias in the Monti consensus clustering method. The results in this manuscript imply a more rigorous approach is required.

## Results

### Systematic bias detected in two widely applied consensus clustering methods

Using clusterlab (see Methods for details), we first generated a null dataset where no genuine clusters are found (Fig. 1a). Next, we tested the Monti consensus clustering algorithm on this data, the cumulative distribution function (CDF) plot corresponding to the consensus matrices from K = 2 to K = 10 for the null dataset demonstrates that as K increases the consensus matrices inherently become more stable (indicated by a flatter line) (Fig. 1b). The PAC scores, which measure the CDF plot flatness, steadily decreased with increasing K estimating an optimal K of ten (Fig. 1c). A similar but reversed effect was observed in the cophenetic metric of Nonnegative Matrix Factorisation (NMF) consensus clustering^15^, which estimates an optimal K of two (Fig. 1d). Therefore, consensus clustering and NMF consensus clustering show bias towards higher and lower values of K, respectively. Both methods also declare evidence of structure when it does not exist, due to not comparing against null reference distributions. To demonstrate the functionality of clusterlab, we generated a ring of four Gaussian clusters, four clusters with varying variance, and a more complex multi-ringed structure consisting of 25 Gaussian clusters (Supplementary Fig. 1).

**Figure 1.**
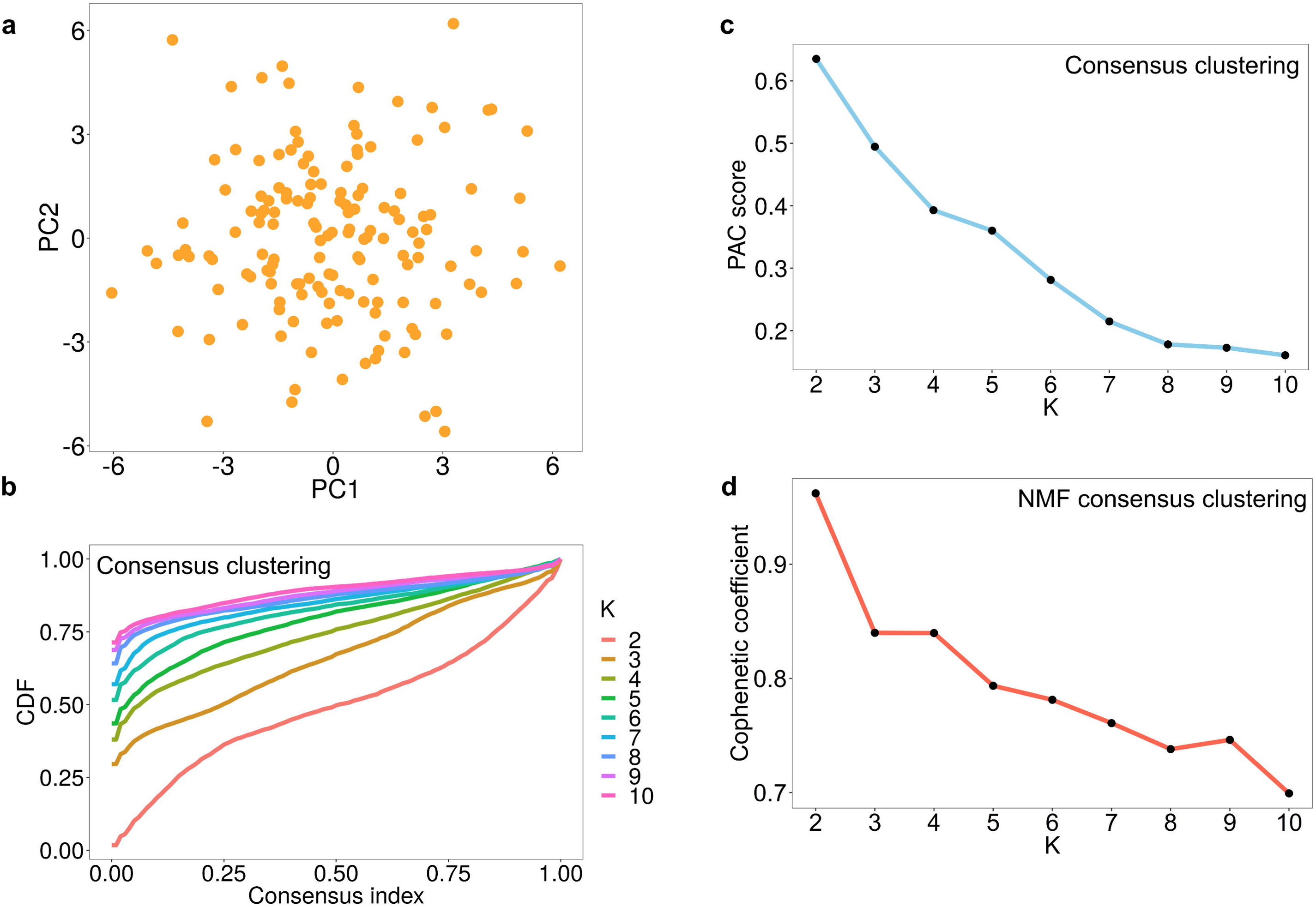
Bias in the estimation of K using Monti and NMF consensus clustering. (A) A PCA plot of a simulated null dataset where only one cluster should be declared. (B) Monti consensus clustering yields a CDF plot implying improved stability with increased K. (C) The PAC score to measure the stability of K decreases with its value, demonstrating a strong preference towards estimating higher optimal values of K. (C) NMF consensus clustering yields a cophenetic coefficient plot which implies lower values of K are preferable using this method.

### M3C can find K and evaluate the significance of its decision

We provide an overview of our method in Figure 2a. For our initial investigations, we tested M3C on a negative control, a simulated dataset in which K = 1 (Fig. 2b). The Relative Cluster Stability Index (RCSI) could not distinguish real from false structure. In contrast, the calculation of Monte Carlo p values by M3C correctly suggested there was no structure in this negative control dataset (alpha = 0.05), and no bias towards higher values of K was observed. Next, M3C was tested on a positive control dataset with four simulated clusters (Fig. 2c). The PAC score and the RCSI correctly identified four as the optimal value of K. A very low Monte Carlo p-value was found by M3C for K = 4 (p = 9.95×10^−21^), this correctly implies that this is the optimal K and means we can reject the null hypothesis H_0_.

**Figure 2.**
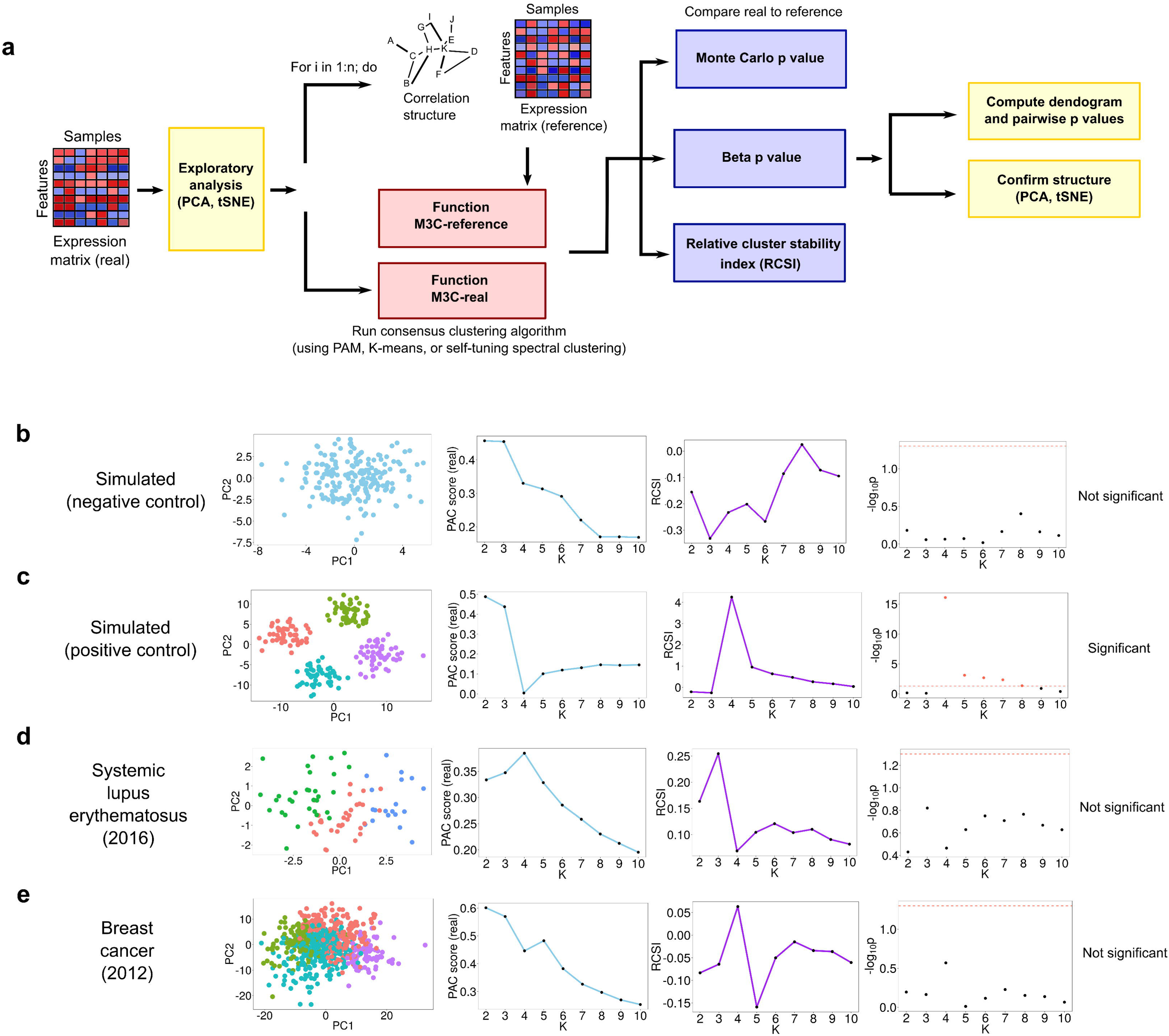
Overview of the M3C method and an initial demonstration. (A) A schematic of the M3C method and software. After exploratory PCA to investigate structure, the M3C function may be run which includes two functions; M3C-ref and M3C-real. The M3C-ref function runs consensus clustering with simulated random data sets that maintain the same gene-gene correlation structure of the input data. While, the M3C-real function runs the same algorithm for the input data. Afterwards, the relative cluster stability index (RCSI), Monte Carlo p values, and beta p values are calculated. Structural relationships are then analysed using hierarchical clustering of the consensus cluster medoids with SigClust to calculate significance of the dendrogram branch points. (B) Results from running M3C on a simulated null dataset, it can be clearly seen that the p values do not reach significance along the range of K, therefore the correct result is suggested, K=1. (C) Results from running M3C on a simulated dataset where four clusters are found, the correct decision is made by M3C. (D) Using M3C, a systemic lupus erythematosus dataset was detected with no significant evidence of structure. (E) Similarly, a breast cancer dataset was identified with no significant evidence of structure.

Next, we reanalysed a range of high-profile stratified medicine datasets where structure had been declared to test for false positive structures (Table 1 & Supplementary Table 1). Because of the ease of data availability, these were predominately, but not exclusively, from TCGA. Table 1 demonstrates the pervasive use of consensus clustering and NMF consensus clustering in the field. Using M3C, we identified two datasets in which no significant evidence against the null hypothesis could be detected. First, a systemic lupus erythematosus (SLE) microarray dataset was analysed where seven major subtypes were reported using hierarchical clustering and dendrogram cutting. However, none of the p-values along the range of K calculated by M3C reached statistical significance (the lowest was for K = 3, p = 0.15) (Fig. 2d). Second, a breast cancer miRNA-seq dataset was identified with no significant evidence of structure (the lowest p value was for K = 4, p = 0.27), whereas seven subtypes were originally reported using NMF (Fig. 2e). These findings imply that false positive structures exist in the literature through not comparing against reference datasets.

**Table I:** Datasets selected for assessment using M3C and optimal K decisions. HC refers to hierarchical clustering and CC to Monti consensus clustering.

| Publication | Year | Data type | Original algorithm | Original K | M3C K |
| --- | --- | --- | --- | --- | --- |
| **Glioblastoma**<sup>3</sup> | 2008 | Microarray | CC | 4 | 4 |
| **Ovarian carcinoma**<sup>4</sup> | 2011 | Microarray | NMF | 4 | 5 |
| **Lung cancer**<sup>5</sup> | 2012 | RNA-seq | NMF | 4 | 2 |
| **Breast cancer**<sup>16</sup> | 2012 | miRNA-seq | NMF | 7 | 1 |
| **Diffuse glioma**<sup>1</sup> | 2016 | RNA-seq | CC | 4 | 8 |
| **Lupus**<sup>9</sup> | 2016 | Microarray | HC | 7 | 1 |
| **Pheochromocytoma**<sup>2</sup> | 2017 | RNA-seq | CC | 4 | 6 |

**Table II.**
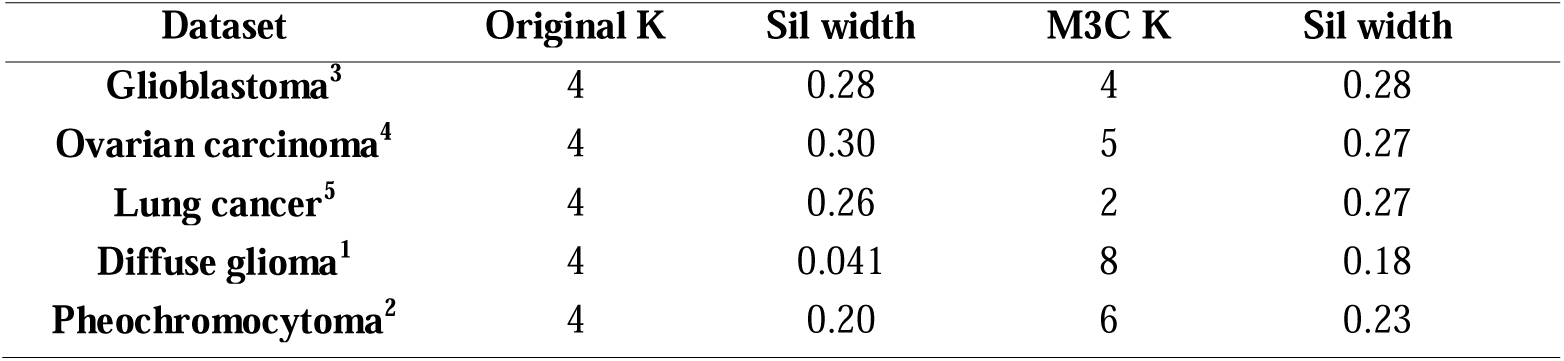
Silhouette width of M3C optimal K assignments compared with original K decision assignments. Higher values of silhouette width (sil width) correspond to preferable clustering.

| Dataset | Original K | Sil width | M3C K | Sil width |
| --- | --- | --- | --- | --- |
| **Glioblastoma**<sup>3</sup> | 4 | 0.28 | 4 | 0.28 |
| **Ovarian carcinoma**<sup>4</sup> | 4 | 0.30 | 5 | 0.27 |
| **Lung cancer**<sup>5</sup> | 4 | 0.26 | 2 | 0.27 |
| **Diffuse glioma**<sup>1</sup> | 4 | 0.041 | 8 | 0.18 |
| **Pheochromocytoma**<sup>2</sup> | 4 | 0.20 | 6 | 0.23 |

### Demonstration of the M3C method on TCGA gene expression data

Of those datasets that exhibited significant evidence of structure using M3C, we used this as an opportunity to contrast the clarity of the M3C results with those from consensus clustering with the PAC score, the NMF cophenetic coefficient^15^, and the GAP-statistic^11^. Our intention in these analyses was not to dispute the original reported K, but instead to test whether methods that do not consider reference distributions along the range of K would lead to visible biases. In these analyses, it was demonstrated that the GAP-statistic continuously increased, implying improving stability regardless of the structure (Supplementary Fig. 2). These findings imply the GAP-statistic is not well suited to analysing complex genome wide expression datasets. Across these datasets, we also demonstrate why M3C fits a beta distribution to the data to estimate extreme tail values, as for K = 2, the beta distribution fits the reference slightly better than a normal distribution (Supplementary Fig. 3 and 4). This step is important as it removes the limitations on p-value derivation imposed by a finite number of simulations (Supplementary Fig. 5).

The PAC score displayed the same bias towards higher K values observed earlier on simulated null datasets, decreasing steadily regardless of the structure, implying increased stability (Figure 3a-e). This effect is more of a problem in datasets where the clustering is not very clear. For the GBM dataset^3^, while a PAC elbow can be seen at K = 4, the global optimal value is K = 10 (Fig. 3a). The problem with the PAC score resembles the problem encountered by Tibshirani, et al. (2001), when the authors developed the GAP-statistic to overcome the subjective decision regarding the location of the elbow. For the GBM case, the Monte Carlo p-values and the RCSI demonstrate a clear optimal value of K = 4 (p = 0.00059), with additional evidence for structure at K = 5 (p = 0.0071).

**Figure 3.**
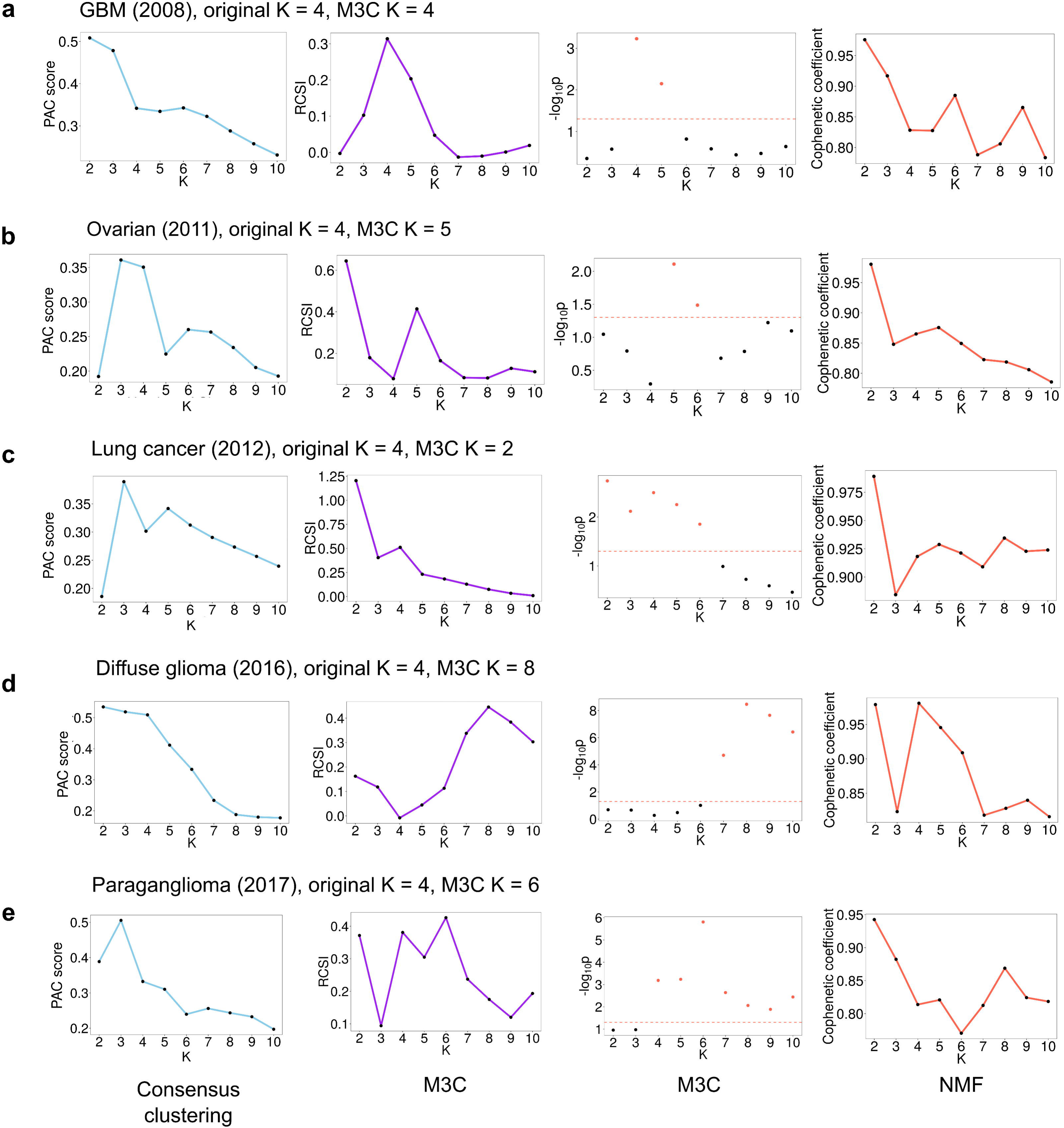
Further evidence of bias existing in widely applied consensus clustering algorithms. (A) Results from running M3C on a glioblastoma dataset^3^ found the optimal K was four. Consensus clustering using the PAC-score shows an optimal K of ten, and NMF of two. (B) Results from running M3C on an ovarian cancer dataset^4^ found the optimal K was five. Consensus clustering using the PAC-score shows an optimal K of two, and NMF also of two. (C) Results from running M3C on a lung cancer dataset^32^ found the optimal K was two. Consensus clustering using the PAC-score shows an optimal K of two, and NMF also of two. (D) Results from running M3C on a diffuse glioma dataset^1^ found the optimal K was eight. Consensus clustering using the PAC-score shows an optimal K of ten, and NMF of four. (E) Results from running M3C on a paraganglioma dataset^2^ found the optimal K was six. Consensus clustering using the PAC-score shows an optimal K of ten, and NMF of two. It can be observed, consensus clustering using the PAC-score and NMF both tend towards K=10 or K=2, respectively, on real data.

For the ovarian dataset^4^, a global optimal PAC value is observed at K = 2, which is supported by the RCSI (Fig. 3b). However, when the Monte Carlo p-values are calculated, it is in fact K = 5 which is the optimal K (p = 0.0078). This happens because some datasets have a skewed null distribution at K = 2, resulting in lower PAC scores (Supplementary Fig. 3b). These are inherently favoured by the algorithm, a bias that is unaddressed by the PAC score or the RCSI. Only by calculating p-values for each value of K can we mitigate against these types of systematic biases.

In cases where the clustering is very clear, the PAC score does perform well. In the lung cancer dataset^5^, a global PAC optimal K can be seen at K = 2, which is supported by both the RCSI and the Monte Carlo p-value (p = 0.0018) (Fig. 3c). Although this conflicts with the original decision of K = 4, the M3C p-value for K = 4 was also significant (p = 0.0032), implying this would be another reasonable choice. However, the bias towards high K values of consensus clustering can be observed again on the diffuse glioma dataset^1^ (Fig. 3d). Here the PAC score continuously decreases until it reaches a global optimum at K = 10. However, considering the reference distributions, M3C informs us that K = 8 is the most significant option (p = 3.5×10^−9^), which is also supported by the RCSI score. For the paraganglioma dataset^2^, the RCSI estimates K = 6 and the Monte Carlo p-value supports this conclusion (p = 1.6×10^−6^), while the PAC score continually decreases, giving no clear choice of K (Fig. 3e). This is another example of why the reference distribution matters, as the RCSI method shows a local maximum for K = 2, while the Monte Carlo p-value does not support this. This is due to the uneven shape of the PG reference distribution for K = 2, which has positive kurtosis (Supplementary Fig. 4b). These findings imply results relying just on relative scores or mean comparisons with the reference can be potentially misleading.

In agreement with our findings on simulated null data, it was observed that the NMF cophenetic coefficient has a tendency towards calling K = 2 on real data (Fig. 3a-e). Only in the diffuse glioma dataset^1^ did the maximum cophenetic coefficient suggest any other value of K. Although there are numerous variant decision rules for NMF in use^4,5,16^, these do not compare against a null distribution. Instead of taking the most stable consensus matrix (highest cophenetic coefficient) as the optimal K, local maxima are often selected^4,5^. Notably, for the ovarian dataset^4^ a local maximum in the NMF cophenetic coefficient was observed at K = 5, which was supported by the M3C decision in this instance. Additional support was observed for the lung cancer optimal K, as an NMF global maximum cophenetic coefficient was detected for K = 2, and the M3C p-value also declared this K to be optimal (p = 0.0018). However, since a tendency in NMF towards K = 2 on null datasets has been observed in this study, it is unclear how confident we should be in this decision.

As a final step, we performed t-Distributed Stochastic Neighbor Embedding (t-SNE) on each dataset then calculated the silhouette width using either the original K or the M3C K to evaluate the relative strength of the M3C cluster assignments. t-SNE was performed first to reduce dimensionality, because the silhouette width has been shown to work poorly alone on high dimensional data in finding the true K^14^. This analysis demonstrated of the four datasets with differing K decisions to the original, the M3C decisions were better in three. These findings support the value of M3C’s reference-based approach to deciding K.

### M3C demonstrates good performance in finding K on simulated data

Next, we sought to evaluate the performance of M3C on simulated data from K = 2 to K = 6 and compare its performance to existing algorithms. In these tests, we varied the clusterlab alpha parameter, which controls the distance between the clusters, and used algorithms which were able to detect the true K from further apart cluster conditions (alpha = 2) to closer ones (alpha = 1) (Fig. 4a,b). Typically, in genome wide analyses many clusters will be overlapping and hard to distinguish from one another. Therefore, sensitivity under these conditions is very valuable. This analysis found that M3C using the RCSI score performed better than consensus clustering with the PAC score, M3C using p-values, the GAP-statistic, CLEST, the original consensus clustering with the delta K score, NMF, and progeny clustering. Notably, while M3C with the RCSI score was approximately 10% higher in accuracy than M3C with p-values, the GAP-statistic, and consensus clustering with PAC, these three methods performed similarly, within 4% of one another. CLEST was also a good performer in this analysis. Overall, these simulations reinforce our findings on real data that M3C performs better than other state-of-the-art methods.

**Figure 4.**
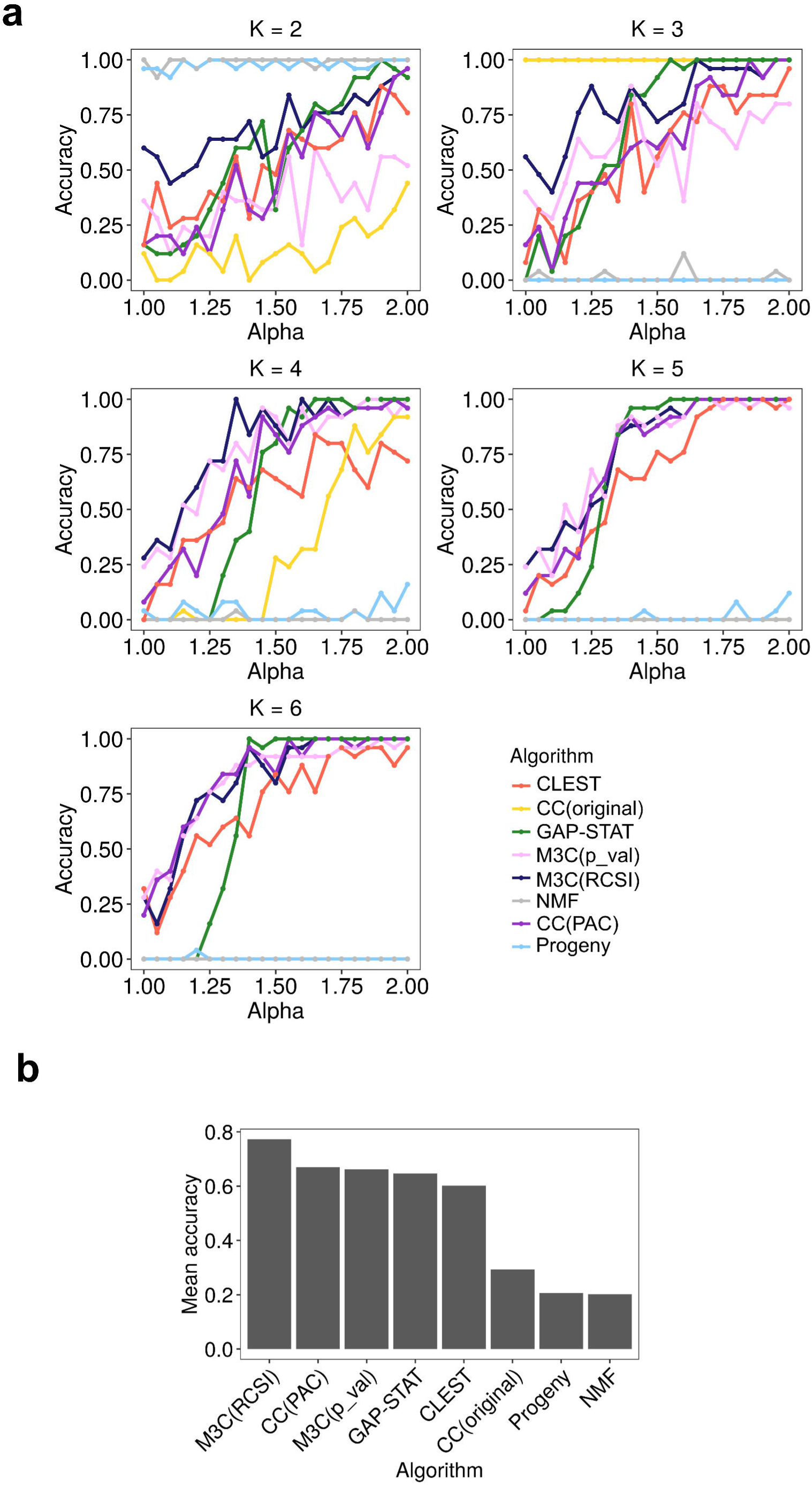
M3C demonstrates good performance in finding K on simulated data. (A) A sensitivity analysis was conducted for every algorithm for K=2 to K=6 while varying the alpha parameter of clusterlab (degree of Gaussian cluster separation). Accuracy was calculated as the fraction of correct optimal K decisions, and for each alpha, with 25 iterations performed at each step. CC(original) refers to the Monti et al. (2003) consensus clustering method, GAP-STAT refer to the GAP-statistic, CC(PAC) refers to consensus clustering with the PAC-score. (B) Performance was calculated across the range of K tested for each algorithm as the mean accuracy.

### M3C can deal with complex structures using spectral clustering

The performance of M3C is dependent on underlying clustering algorithm. Although k-means and PAM perform well on the types of data generally encountered in genome-wide studies, they assume the clusters are approximately spherical and equal in variance, which may not be true. Spectral clustering is a widely applied technique due to its ability to cope with a broad range of structures^17^. Therefore, to increase the capabilities of the M3C software package, it includes self-tuning spectral clustering^18^. We tested spectral clustering as M3C’s inner algorithm versus PAM and k-means on two synthetic datasets, one where the clusters were anisotropic (Fig. 5a), and a second where one cluster had a far smaller variance than its neighbouring cluster (Fig. 5b). Under these conditions, it was observed that M3C using PAM and k-means both had problems identifying the true K and classifying the members of each cluster correctly. On the other hand, M3C using spectral clustering did not suffer these drawbacks. Using spectral clustering, M3C is also capable of recognising more complex non-Gaussian shapes, such as half-moons and concentric circles (Supplementary Fig. 6). The addition of spectral clustering to the M3C software package allows greater flexibility in the range of structures that may be examined.

**Figure 5.**
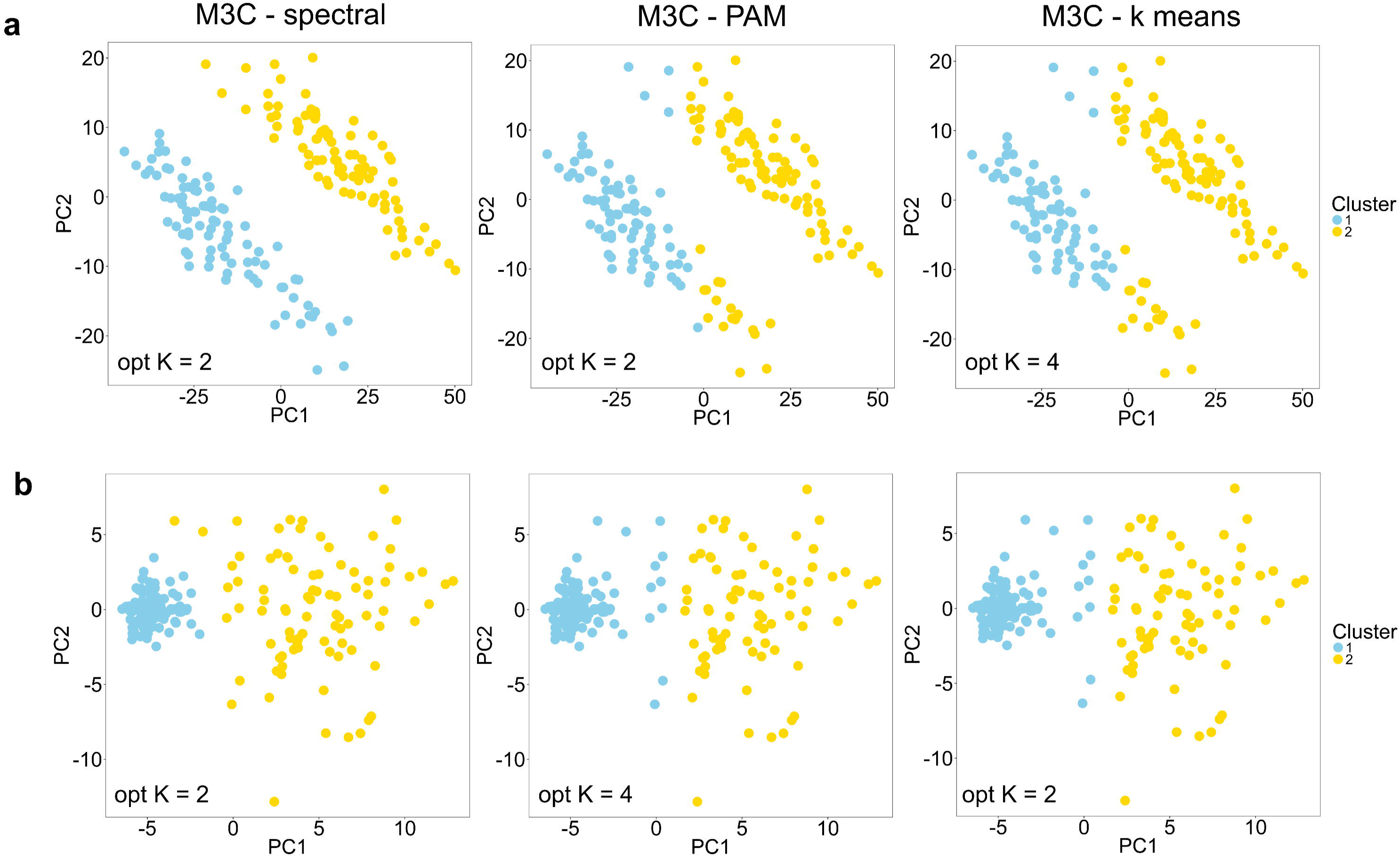
M3C uses spectral clustering to deal with complex structures. (A) Results from running M3C using either spectral, PAM, or k-means clustering on anisotropic structures. The results for K=2 for each inner algorithm are shown in all cases, in the corner of the plots are the optimal K decisions using the RCSI. (B) Similarly, results from testing different internal algorithms on structures of unequal variance.

### M3C can quantify structural relationships between consensus clusters

An important question when the optimal K has been decided is, how do the discovered clusters relate to one another? Inherently, consensus clustering does not distinguish between flat versus hierarchical structure. To solve this, M3C performs hierarchical clustering on the medoids of each consensus cluster. To make the analysis statistically principled, M3C iteratively performs the SigClust method^19^ on each pair of consensus clusters, then displays the pairwise p-values for each split of the dendrogram. Testing M3C on the PG dataset revealed a hierarchical relationship between the six clusters (Fig. 6a), with, for example, consensus clusters one and two grouping together (p = 1.2×10^−80^). In contrast, testing M3C on a null dataset without clusters demonstrated insignificant SigClust p-values and a flat dendrogram (Fig. 6b). The addition of a hierarchical clustering stage after choosing the optimal K should prove helpful in identifying structural relationships.

**Figure 6.**
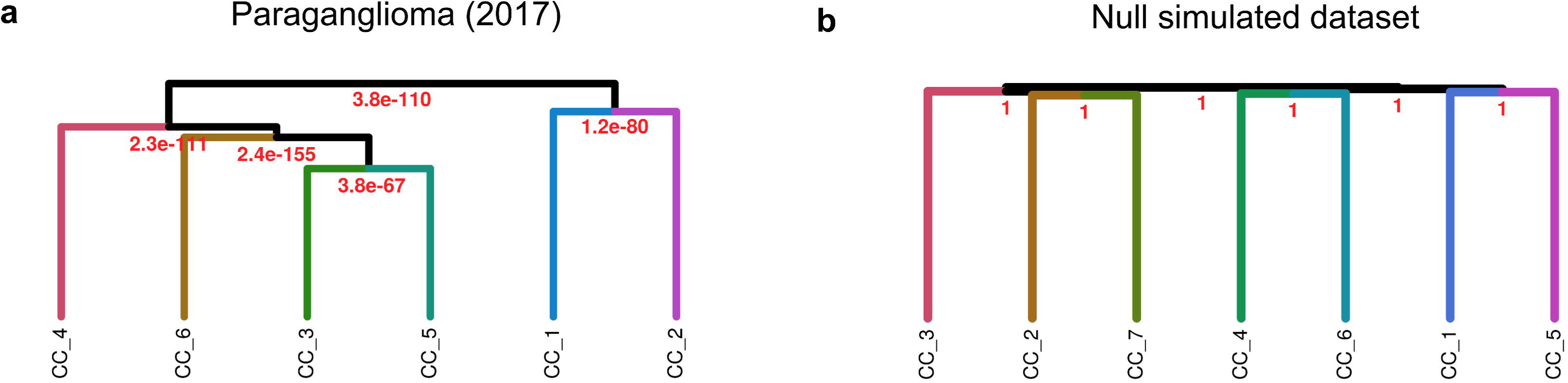
M3C can investigate structural relationships between consensus clusters. M3C calculates the medoids of each consensus cluster, then hierarchical clustering is performed on these, SigClust is run to detect the significance of each branch point. (A) Results from M3C structural analysis of the six clusters obtained from the paraganglioma dataset analysis^2^, all p values were strongly significant, supporting the M3C decision of the declaration of structure. (B) Results from the same analysis run on a simulated null dataset of the same dimensions, no p values were significant.

### Sensitivity and complexity analysis of M3C

As a final step, we decided to evaluate M3C’s internal parameters using the PAM algorithm, compare its runtimes with other methods, and calculate its complexity. A sensitivity analysis of the number of inner replications and outer simulations found M3C generally yielded stable results across six TCGA datasets with 100 inner replications and 100 outer simulations (Supplementary Fig. 7-8). We executed M3C on five datasets on a high-powered desktop computer using a single thread of an Intel i7-5960X CPU @ 3.00GHz with 32GB of RAM. Runtimes ranged between 2-25 minutes, depending on dimensionality (Fig. 7a). We compared the runtime of M3C with other well performing methods from our earlier analysis on the same computer with a single thread (Fig. 7b-c). M3C, CLEST, and the GAP-statistic which all use Monte Carlo simulations as a reference were set to 25 reference iterations for comparative purposes. This analysis demonstrated that consensus clustering with the PAC score was the fastest method, followed by the GAP-statistic. CLEST and M3C were slower and similar in runtime for lower N (number of samples), but for N greater than 500, M3C performed more slowly than CLEST (Fig. 7b).

**Figure 7.**
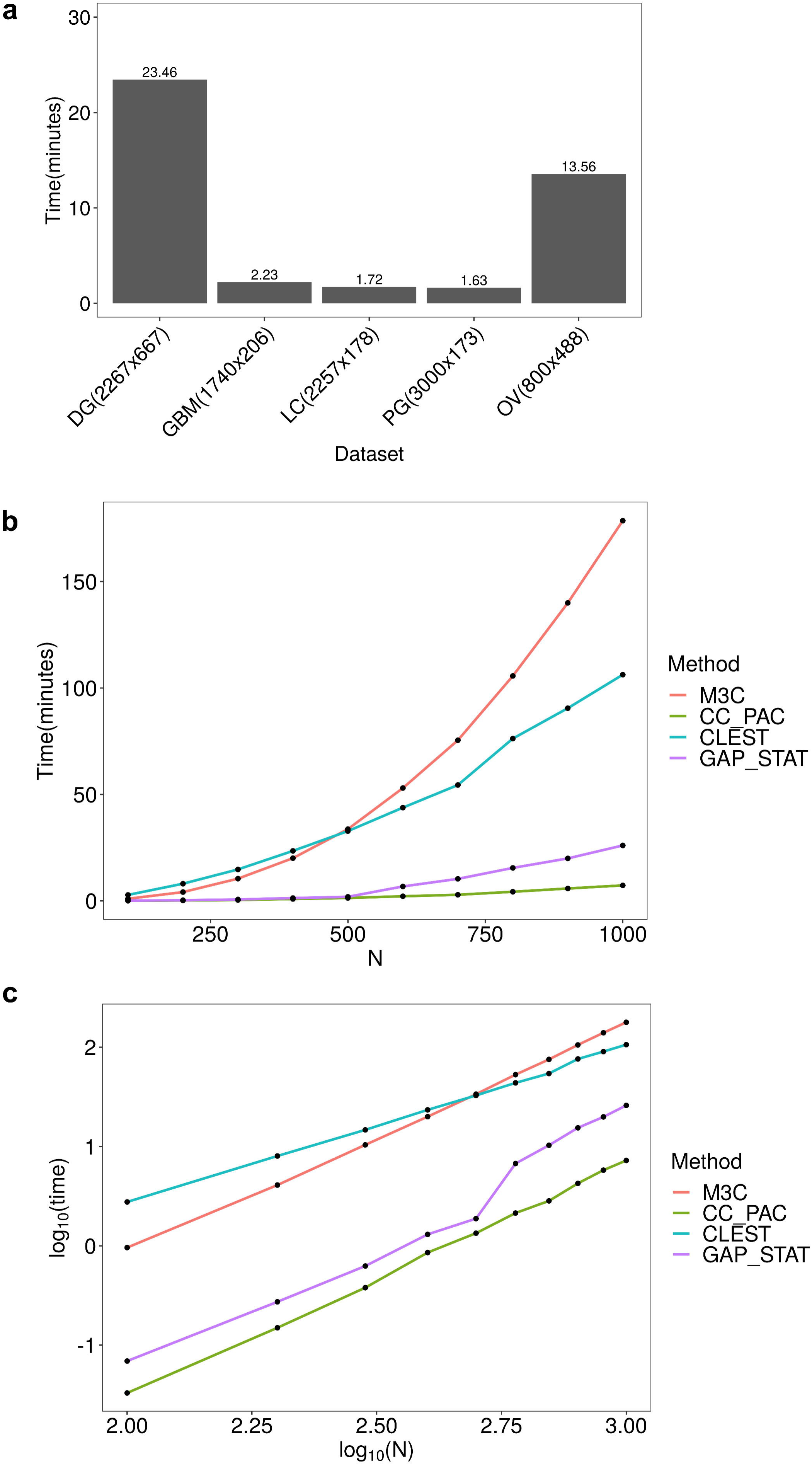
M3C can perform quickly across a range of datasets. (A) M3C runtimes (in minutes) for five datasets used in the analysis. Performance was measured on an Intel Core i7-5960X CPU running at 3.00GHz using a single thread with 32GB of RAM. M3C was run using 25 outer Monte Carlo simulations and 100 inner iterations using the PAM algorithm. (B) M3C and other method runtimes in minutes for a series of simulated datasets with the number of samples (N) ranging from 100-1000 for datasets of 1000 features. CLEST and the GAP-statistic, which also use a Monte Carlo reference procedure, were set to run with 25 Monte Carlo simulations, the same as M3C for comparison. (C) Log-log plot of the same data shown in B.

The complexity of the M3C algorithm is *O*(*BHA*/*C*), where *B* is the number of Monte Carlo simulations, *H* is the number of consensus clustering resamples, and *A* is the complexity of the underlying clustering algorithm (see pseudo-code for M3C in Supplementary Note 1). The *C* denotes number of available processors, as M3C can be parallelized due to its independent simulations and subsampling subroutines. We empirically evaluated M3C’s time complexity as a function of sample size *N* using the PAM algorithm, which has a complexity of *O*(*N*^2^). Calculating the slope of the log-log plot yielded an empirical complexity of *O*(*N*^2.4^). This demonstrates that M3C is approximately quadratic in *N*.

## Discussion

We report the advancement of the Monti consensus clustering algorithm to include a Monte Carlo simulation driven reference system for estimating the optimal K and testing the null hypothesis K=1, we call the method M3C. Our investigation into this consensus clustering algorithm demonstrated it has inherent bias towards higher values of K. These occur due to not considering the reference distribution along the range of K when deciding on its value. Although considering these distributions is a relatively straightforward procedure, as we have demonstrated, it has important implications. To date, testing of the null hypothesis by TCGA has been conducted by SigClust after deciding on the value of K using the standard methods^2,6,16^. SigClust tests the null hypothesis K=1 for pairs of clusters, but it does not directly estimate K. The advantage of M3C is that it can both find K and test the null hypothesis K=1.

Our reanalysis of high-profile stratified medicine studies, predominantly from TCGA^1-5,9,16^, questions the value of consensus clustering when used without considering the appropriate reference distributions. The bias towards higher values of K, coupled with subjective decision making as to what constitutes the optimal K, similar to the original elbow problem solved by the GAP-statistic^11^, may provide misleading results. We identified two cases in the literature where structure had been declared despite M3C indicating no significant evidence against the null hypothesis. In the case of the SLE study, seven subtypes were originally declared in a major transcriptomic analysis^9^. Within the context of these new findings, it is perhaps better to describe these subtypes as existing within a noisy spectrum of non-distinct states. This hints that there may be publication bias for positive declaration of structures.

It is necessary to remark on the limitations of the approach. The M3C method can allow testing of the null hypothesis K=1 and mitigate bias. However, this method does not allow, for example, the formal statistical comparison of selecting K=2 compared with other values of K. The relative magnitude of the p values can be used to estimate the optimal K by comparing against the null K=1 scenario like using the RCSI, however, this is not formal hypothesis testing. A second limitation is that M3C is computationally expensive, however, extreme tail estimation and multi-core ability mitigate this problem. Finally, just because the p-value or RCSI supports a given K gives no guarantee the identified clusters or their number will be reproducible in an independent validation dataset.

Other types of consensus clustering methods include Infinite Ensemble Clustering^20^ (IEC) and Entropy-based consensus clustering^21^ (ECC), which can be used for patient stratification. IEC incorporates marginalized denoising auto-encoder with dropout noises to generate the expectation representation for infinite basic partitions. ECC employs an entropy-based utility function to fuse many basic partitions into a single consensus structure. A future challenge is to systematically evaluate the performance of a wider range of consensus clustering methods on genome wide expression data.

We benchmarked the performance of M3C against a number of alternatives, including: Monti consensus clustering, the GAP-statistic, progeny clustering, and CLEST. Several cluster validity indices were not tested, however, such as: the Silhouette index^22^, the Calinski Harabasz index^23^, the Jaccard index^24^, and the Davies-Bouldin index^25^. It would be interesting to determine if any of these indices perform well in determining the optimal K when applied on consensus matrices produced by the consensus clustering algorithm, our study indicates they will be subject to bias without a reference procedure. It is also relevant to mention that there are other methods that could be applied to investigate the significance of dendrogram splits, such as the inheritance procedure^26^.

Lastly, it is important to mention the methodological contributions of clusterlab. Clusterlab is a flexible new method for generating Gaussian clusters. Unlike prior methods^14,27,28^, it is able to generate and position Gaussain clusters in a highly customisable manner with specified variance, spacing, and size. Clusterlab can generate data similar in nature to cancer gene expression datasets, which are typically high-dimensional and Gaussian^19^. The method should appeal to researchers in a range of disciplines for testing methods for finding K and clustering algorithms.

## Methods

### M3C

The method uses a Monte Carlo simulation, which generates random data with each iteration, to repeat the Monti et al. consensus clustering algorithm many times over. Then, the real algorithm is run just once to compare the real cluster stabilities along the range of K with those expected using random Gaussian data (K = 1). Pseudo-code is given in Supplementary Note 1. This gives a new method for choosing K after consensus clustering that removes bias towards high values of K and allows one to statistically test for the presence of structure. The specific details are now given.

#### Simulation of the reference dataset

There are a range of options for the generation of reference datasets in M3C’s Monte Carlo simulation. We use an approach first proposed by Tibshirani et al., which preserves covariance structure via principal component analysis (PCA). With an input matrix, *T* ∈ ℝ^*S*\**F*^ we can compute the input data’s eigenvector matrix *A* ∈ ℝ^*F*\**S*^ and its principal component score matrix, *Y* ∈ ℝ^*S*\**S*^, where *F* is the number of features in the provided matrix, and *S* is the number of samples. The steps taken to generate random data are repeated *b*= 1 … *B* times:

1. Conduct PCA to obtain the orthogonal matrix of eigenvectors, *A* of the input data *T*:

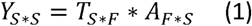
2. Next, a random PC score matrix is generated, *Y*^*b*^ ∈ ℝ^*S*\**S*^, where the th column is filled with random values from a normal distribution with mean zero and standard deviation equal to the *i*th column in *Y*. Let, *D*_*i*_ be the standard deviation of *Y*_\**i*_ and for *I* = 1 … *S*:

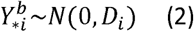
3. Multiplying *Y*^*b*^ with the transpose of *A* yields *Q*^*b*^ ∈ ℝ^*S*\**F*^, a single simulated null dataset with the same feature correlation structure as *T*, but without clusters.

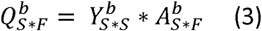

Steps 1-3 are repeated by M3C for each Monte Carlo reference simulation for *b* = 1 … *B*, and for the bth simulation one random dataset, *Q*^*b*^ is passed into the consensus clustering algorithm (described below) to calculate null reference stability scores for *K* = 2, …, *maxK*. After *B* simulations, the consensus clustering algorithm is run just once on the input data for comparison using procedures we will go on to detail. M3C is set to use *B* = 100 and this was the parameter setting used for the simulations in this study.

#### Consensus clustering

The Monti et al. consensus clustering algorithm subsamples the input data sample-wise, *H* times, and with each resampling iteration clusters the perturbed dataset using a user defined inner clustering algorithm (e.g., PAM) for each value of *K*. It then measures the stability of the sample cluster assignments over all resampling iterations to decide *K*. M3C includes PAM, k-means, and spectral clustering as options, with PAM set by default due to its superior speed. Let, *D*^(1)^, *D*^(2)^,…, *D*^(*H*)^ be the list of *H* perturbed datasets, and let *M*^(*h*)^ … {0,1} ^*N*\**N*^ be the connectivity matrix resulting from clustering dataset *D*^(*h*)^, the entries of *M*^(*h*)^ are then defined as:

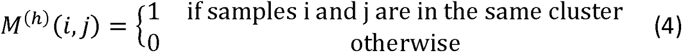

To keep count of the number of times samples *i* and *j* are resampled together in the perturbed dataset *D*^(*h*)^ an indicator matrix *I*^(*h*)^ ∈ {0,1} ^*N*\**N*^ is defined:

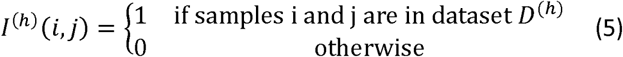

The consensus matrix, *M* ∈ [0,1]^*N*\**N*^, is defined as the normalised sum of all the connectivity matrices of all *H* perturbed datasets:

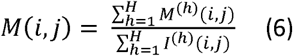

The entry (*i,j*), or consensus index, is the number of times that two samples cluster together divided by the total number of times they were sampled together across all the perturbed datasets. A value of 1 would correspond to a perfect score as the two samples are always found in the same cluster across all resampling runs, while a value of 0 would correspond to the worst score as the two samples never are found in the same cluster. A consensus matrix is generated for every value of *K* and then the stability of each matrix quantified using an empirical cumulative distribution (CDF) plot. For any given consensus matrix *M*, the CDF is calculated and is defined over the range [0,1] as follows:

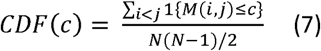

Where 1{…} denotes the indicator function, *M*(*i,j*), denotes entry (*i,j*) of the consensus matrix *M, N* is the number of rows (and columns) of *M*, and *c* is the consensus index value.

#### Calculation of the PAC score

The CDF plot has consensus index values on the x axis and CDF values on the y axis. A perfectly stable cluster solution will have a flat CDF plot representing a matrix purely of 0s and 1s, therefore the degree of CDF flatness for each *K* is a measure of the stability of *K*. To quantify this, M3C uses the PAC score, a metric shown to perform well in simulations^14^. PAC is defined as the fraction of sample pairs with consensus index values falling in the intermediate sub-interval (*x*_1_,*x*_2_) ∈ [0,1]. For a given value of *K, CDF*(*c*) corresponds to the fraction of sample pairs with consensus index values less than or equal to c and PAC is defined as:

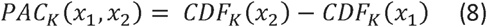

M3C calculates the PAC score with *x*_1_= 0.1 and *x*_2_ = 0.9. Although the PAC window is a user defined parameter, we have found these settings to perform well in our experience.

#### Calculation of the RCSI

To account for the reference PAC scores from *b* = 1 … *B*, where *B* is the total number of Monte Carlo simulations, M3C uses the RCSI. Let, *Pref*_*Kb*_ be the reference PAC score from the *b*th Monte Carlo simulation for a given *K*, and, *Preal*_*K*_ the real PAC score for that *K*, then the *RCSI*_*K*_ is defined as:

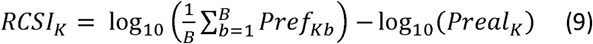

#### Calculation of the Monte Carlo p value

To improve the selection of the optimal *K*, M3C derives Monte Carlo p values by testing the real PAC score for each *K* against the null PAC distribution, generated using simulated structureless data. Let *o*_*K*_ be the number of observed PAC scores in the reference less than or equal to the real PAC score, let *B* be the total number of Monte Carlo simulations, and the p value for that value of K, *P*_*K*_ is then defined as:

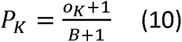

Where 1 is added the numerator and denominator to avoid p values of zero^29^.

#### Interpretation of the p-values

For each *K* the method will test the null hypothesis *H*_0_ that the PAC score, *Preal*_*K*_, came from a single Gaussian cluster (*K* = 1) versus the alternative hypothesis *H*_*A*_ that *Preal*_*K*_ did not come from a single Gaussian cluster (*K* ≠ 1). If a p value for a *K* reaches significance (alpha=0.05) it should be viewed as evidence that the data is not a single Gaussian cluster. If no p values along the range of *K* reaches significance (alpha=0.05) it should be viewed as evidence that the data is a single Gaussian cluster. The relative significance of the p-values can be used to suggest the most preferable *K*, although we caution that the method does not formally test the selection of one value of *K* versus another.

#### Calculation of the beta distribution p-value

To estimate p-values beyond the range of the Monte Carlo simulation, M3C fits a beta distribution. This distribution is more flexible than the normal alternative, which is especially helpful when K = 2, which tends to result in null distributions with nonzero skew and kurtosis. Moreover, the PAC score is bound on the interval [0,1], as is the beta distribution, providing the correct range for computation. The *α* and *β* shape parameters required for the beta distribution are derived using maximum likelihood estimates for the mean, *μ*, and variance, *σ*^2^, of the reference PAC scores for any given K:

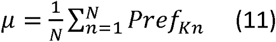

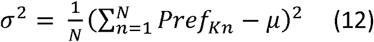

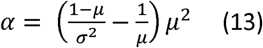

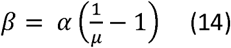

These *α* and *β* shape parameters are then used by M3C to generate the reference distribution for K. The real PAC score is used as a test statistic for derivation of the estimated p value. Let *x* denote the reference PAC score. Then the beta probability density function (PDF) is defined as:

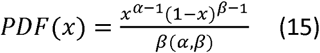

### Simulating NXN dimensional Gaussian clusters in a precise manner

We found that current Gaussian cluster simulation methods were inadequate for systematic testing of M3C. MixSim^27^, generates Gaussian clusters, however, it is not possible to precisely control their positioning. The Python scikit-learn machine learning module contains a Gaussian cluster simulator, but it generates clusters randomly and controlled positioning is not possible. Another method allows controlled spacing^14^, but does not generate Gaussian clusters, instead the clusters resemble triangular slices and the variance and size cannot be set. Therefore, we developed clusterlab (https://cran.r-project.org/web/packages/clusterlab/index.html). Clusterlab is a novel method that allows simulation of Gaussian clusters with controlled spacing, size, and variance. It works by generating cluster centres or points on the circumference of a circle in 2D space because this is easier to work in mathematically than higher dimensional space. The specific details are now given.

#### Generating evenly spaced points on the perimeter of a circle

To control the spacing, size, and variance of synthetic clusters, clusterlab works within a 2D Cartesian coordinate system with an origin at (0,0). First, the algorithm generates a set *S* = {*w*_*i*_ ∈ ℝ^2^, *i* = 1, … *X*} of *X* evenly spaced pairs of coordinates, where *w*_*i*_ = (*x*_*i*_, *y*_*i*_), on the perimeter of a circle. Each of these coordinates later will be the centre of a Gaussian cluster, therefore, *X* is also the number of clusters to be generated. Let, *r* be the radius of the circle, then, for the *i*th cluster centre from *i* = 1 … *X* we need to set *i* = 0 for the first cluster centre, so for *i* = 0 … *X* − 1, the coordinate pairs are calculated as follows:

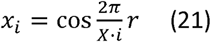

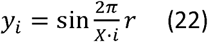

This naturally leaves the *r* parameter as a means of controlling the spacing of the cluster centres. However, at this point, we also introduce an additional parameter for moving the *i*th cluster centre, *α*_*i*_. *α*_*i*_ is a scalar that can be used to push each coordinate pair (or vector) away from its starting point, yielding the transformed coordinates 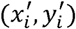. In the case of a cluster being left stationary, *α*_*i*_ = 1. More specifically, for all pairs in set *S*, from *i* = 1… *X*:

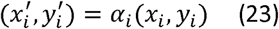

We also leave the option to add a final coordinate to *S* at (0,0), to allow a central cluster within the middle of the ring to be generated later.

#### Generation of more complex multi-ringed structures

As an optional next step to extend the single ring system, clusterlab can create multiple rings or concentric circles of 2D coordinates. After simulating the *q*th ring, as described above, from *q* = 1 … *Q*, the *q*th rings 2D coordinates are pushed away from the origin using vector multiplication with a scalar, let this scalar be *β*_*q*_, let the newly transformed coordinates be 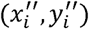, and so for *i* = 1 … *X*:

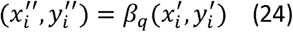

Our new total number of samples, *T*, will be, *T* = *X* * *Q*. With each iteration from *q* = 1 … *Q*, the *i*th transformed coordinates, 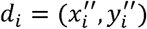, are added to a new set, *R*= {*d*_*i*_ ∈ ℝ^2^, *i* = 1,… *T*}. Optionally, another layer of complexity may be added by using vector rotations of the *q*th rings coordinate pairs from *i* = 1 … *X*, by setting *θ*_*q*_ ≠ 0 in the following equation. To calculate each of the rings new coordinates 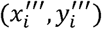 from *i* = 1 … *X*, the following calculation is performed for every pair:

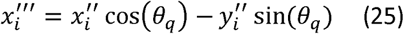

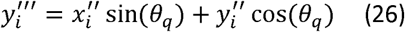

#### Generation of Gaussian clusters

At this point we will assume that multiple rings have not been generated and we are working with, *S*, a set of 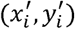 coordinates described by equation 23. However, the method that generates the Gaussian cluster multi-ringed system is identical to the single ringed system described below, except we start with the multiplied 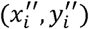 or multiplied and rotated set of 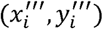 points from the multi ring 2D coordinate set, *R*.

To form *X* Gaussian clusters of size *M*_*i*_ per cluster, we add Gaussian noise from a normal distribution, *N*(0,*D*_*i*_), to the *i*th pair of cluster centre 2D coordinates, 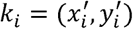, to create the new coordinates, *t*_*i*_ = (*x*_*j*_,*y*_*j*_). Performing this *M*_*i*_ times for each cluster centre, giving a total of 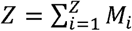 coordinate pairs, yields the final set, *J* = {*t*_*i*_ ∈ ℝ^2^, *i* = 1, … *Z*}. The number of samples in each cluster may be set by varying *M*_*i*_, and the clusters variance, by setting *D*_*i*_. The new coordinate pairs, (*x*_*j*_, *y*_*j*_), to be added to, *J*, for all samples are calculated as follows:

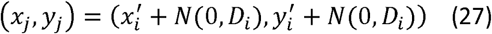

#### Projection of the final 2D coordinates into N dimensions

We transform the cluster sample coordinates into *N* dimensions with a previously explained method which uses a reverse PCA^14^. First, two random vectors are generated of length *V*, where *V* will equal the number of features in the final matrix, from a normal distribution *N*(0,0.1), let these be *v*_1_and *v*_2_. The SD of 0.1 was chosen empirically after examination of the scale of the simulated PC plots compared to those from real expression datasets. The *v*_1_ and *v*_2_ vectors are treated as fixed eigenvectors in this method, and each of our previously simulated coordinate pairs are treated as 2D PC scores. The final matrix, *F* ∈ ℝ^*Z*\**V*^, comprised of *Z* rows (samples) and *V* columns (features), is formed by linear combinations of the fixed eigenvectors with the pairs of PC scores. Let, *x*_*i*_ and *y*_*i*_ be the PC scores of the *i*th sample, from *i* = 1 … *Z* from set *J*, then the *i*th row of the output matrix *F* is given by:

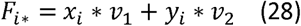

#### Non-Gaussian structures

For generating structures used in the spectral clustering analysis, the CRAN clusterSim package version 0.47 was used^30^. For the anisotropic and unequal variance clusters, 90 samples were simulated with two dimensions with the cluster.Gen function using the default settings. For the half-moon clusters, the shapes.two.moon function was used with 90 samples, and for the concentric circles the shapes.two.circles function was used with 180 samples, both using default settings. The sample number was increased in the latter to prevent gaps forming in the concentric circles.

### Real test datasets

All test datasets, apart from the SLE dataset, were already normalised and downloaded directly through TCGA publication page (https://tcga-data.nci.nih.gov/docs/publications/) during the period of April to June 2017, further details are provided in Supplementary Table 1. We chose RNA-seq or microarray data from the TCGA where the data was already normalised. The diffuse glioma (DG) dataset is a RNA-seq matrix consisting of 2266 features and 667 samples^1^ (https://tcga-data.nci.nih.gov/docs/publications/lgggbm_2015/LGG-GBM.gene_expression.normalized.txt). The GBM dataset, is a microarray matrix consisting of 1740 features and 206 samples^3^ (https://tcga-data.nci.nih.gov/docs/publications/gbm_exp/unifiedScaledFiltered.txt), the feature list used was taken from a later publication on the same dataset^6^. The lung cancer (LC) dataset^5^ used was a RNA-seq matrix consisting of 178 samples and 2257 features (https://tcga-data.nci.nih.gov/docs/publications/lusc_2012/gaf.gene.rpkm.20111213.csv.zip), the feature list used to filter this dataset was from an earlier publication where four subtypes had been identified (http://cancer.unc.edu/nhayes/publications/scc/wilkerson.scc.tgz). The paraganglioma (PG) dataset downloaded was a RNA-seq matrix consisting of 173 samples and 3000 features (https://tcga-data.nci.nih.gov/docs/publications/pcpg_2017/PCPG_mRNA_expression_naRM.log2.csv.zip), the gene wise filtering scheme used was the same as described as in the corresponding publication^2^. The ovarian cancer (OV) dataset^4^ was a RNA-seq matrix of 489 samples and 800 features (https://tcga-data.nci.nih.gov/docs/publications/ov_2011/TCGA_489_UE.zip), and the gene list used for subsequent filtering was obtained from an earlier publication that detected four subtypes^31^. The SLE dataset^9^ used was a microarray matrix of 82 samples and 48 features, the data was obtained from GEO (GSE65391), normalised, and filtered in the manner described in the associated publication.

## Author contributions

C.R.J conceived and designed the approach. C.R.J and D.W wrote the manuscript. C.R.J and D.W wrote the code. C.R.J, D.W, K.G, and D.R performed data analyses. All authors reviewed and edited the manuscript. M.B, C.P, M.E, and M.L supervised the project.

## Additional information

The authors declare that they have no competing interests.

