## Supplementary Information for "M3C: Monte Carlo reference-based consensus clustering"

**Supplementary figure 1**

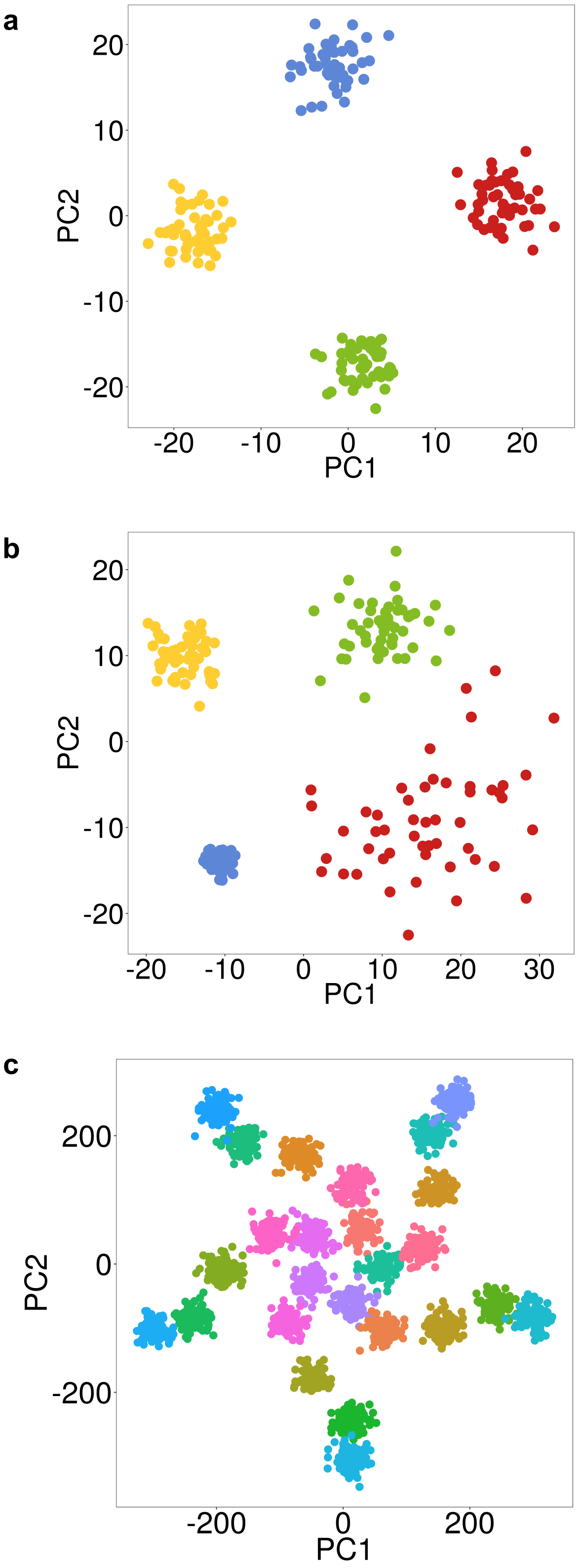

**Supplementary figure 1.** Clusterlab demonstration of the simulation of three synthetic datasets. For explanation of parameters see methods. (A) Simulation of 4 Gaussian clusters with $D = 1$, $M = 50$, $r = 8$. (B) Simulation of 4 Gaussian clusters with $D = (0.5,1,1.5,3)$, $M = 50$, $r = 8$. (C) Simulation of 5 rings of 5 Gaussian clusters with $D = 6$, $M = 100$, $r = 7$, $\beta = (2,4,6,8,10)$, $\theta = (30,90,180,0,0)$.

**Supplementary figure 2**

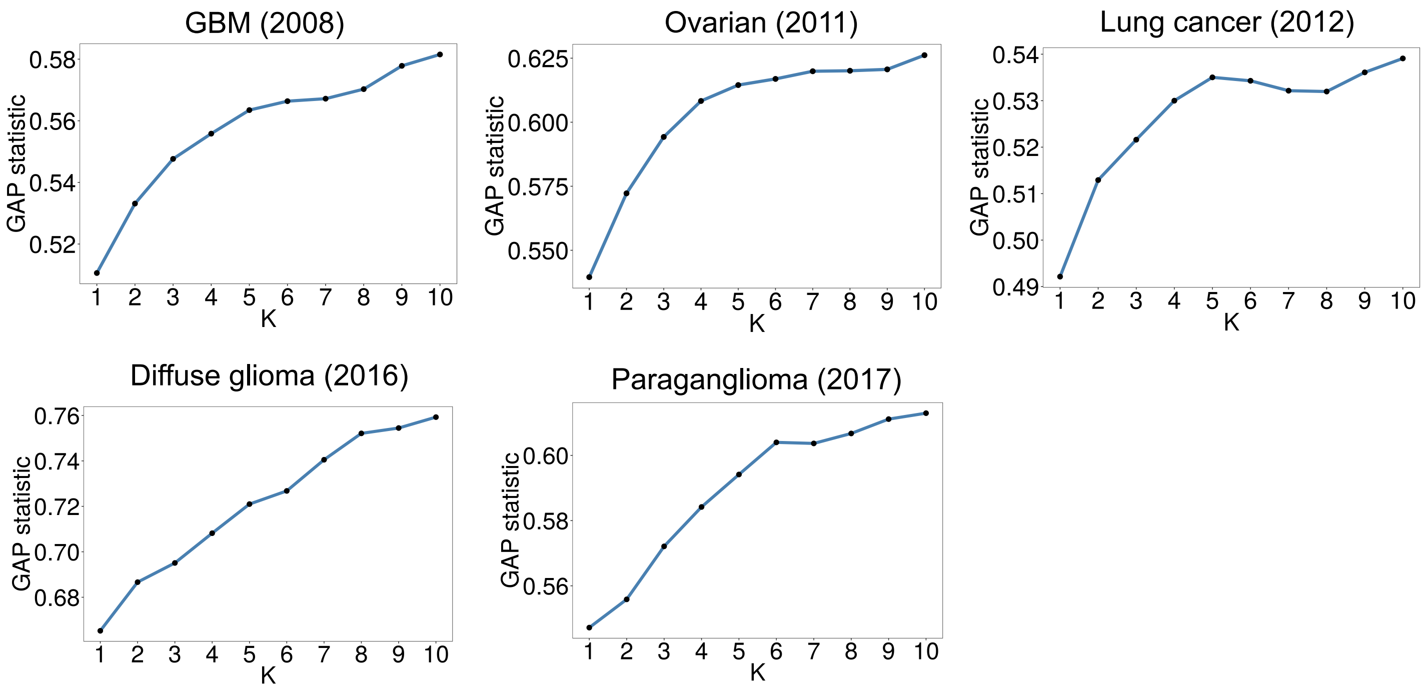

**Supplementary figure 2.** Results from running the GAP-statistic across 5 datasets using the PAM algorithm. A potentially misleading trend towards increased stability at higher values of K can clearly be observed.

**Supplemental figure 3**

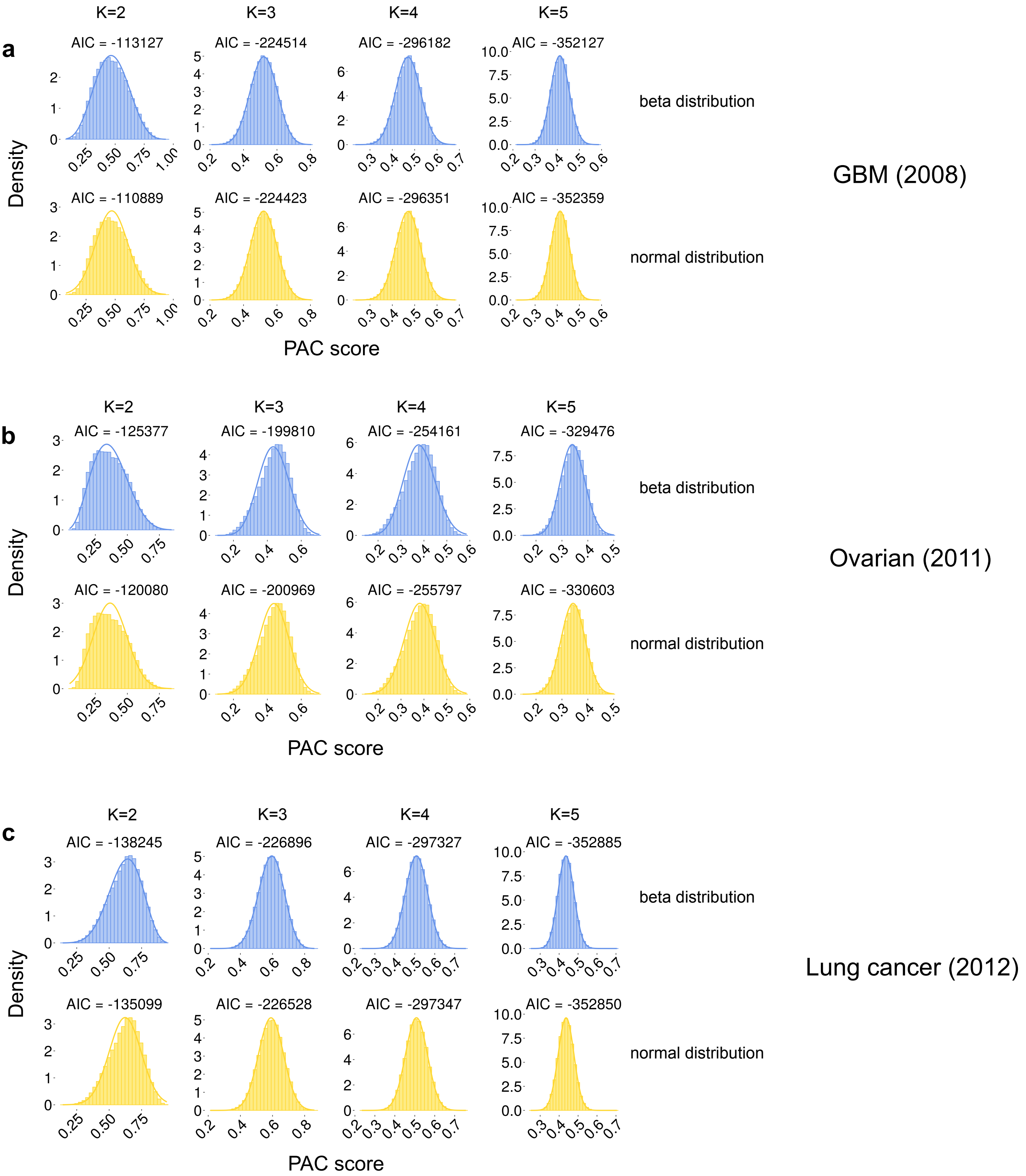

**Supplementary figure 3.** Reference distributions generated using M3C of PAC scores across the range of K from 2-5. 100,000 simulations were performed to generate the distributions and a normal (yellow) and beta (blue) fit were compared. Using the Akaike information criterion (AIC) it can be observed that for K=2, a beta distribution fits slightly better than the normal (indicated by a more negative AIC). This is due to the skew and kurtosis that the reference distributions often have at K=2.

**Supplemental figure 4**

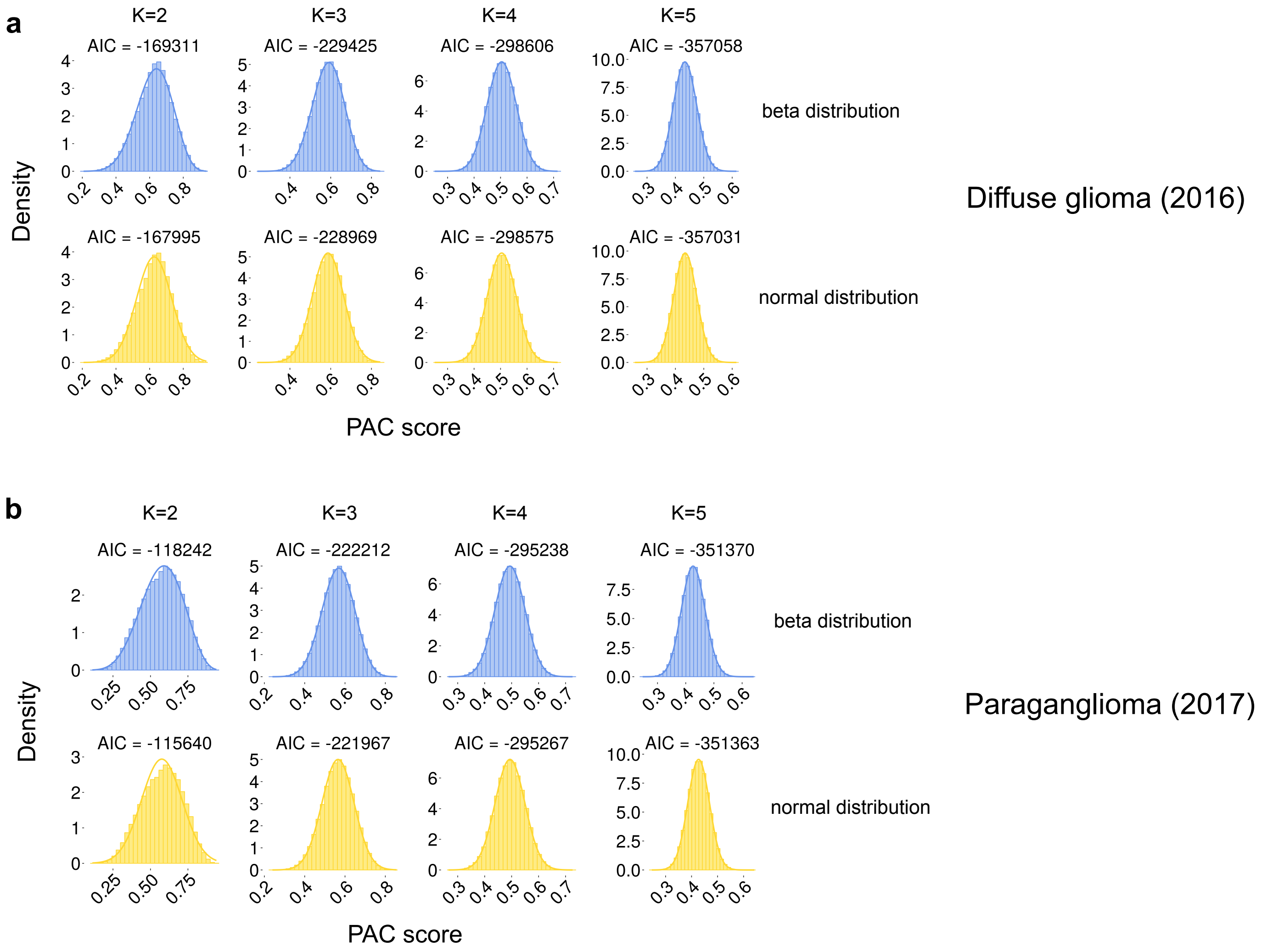

**Supplementary figure 4.** See supplementary figure 3 legend for details, the same analysis was performed, but with different datasets.

### Supplementary figure 5

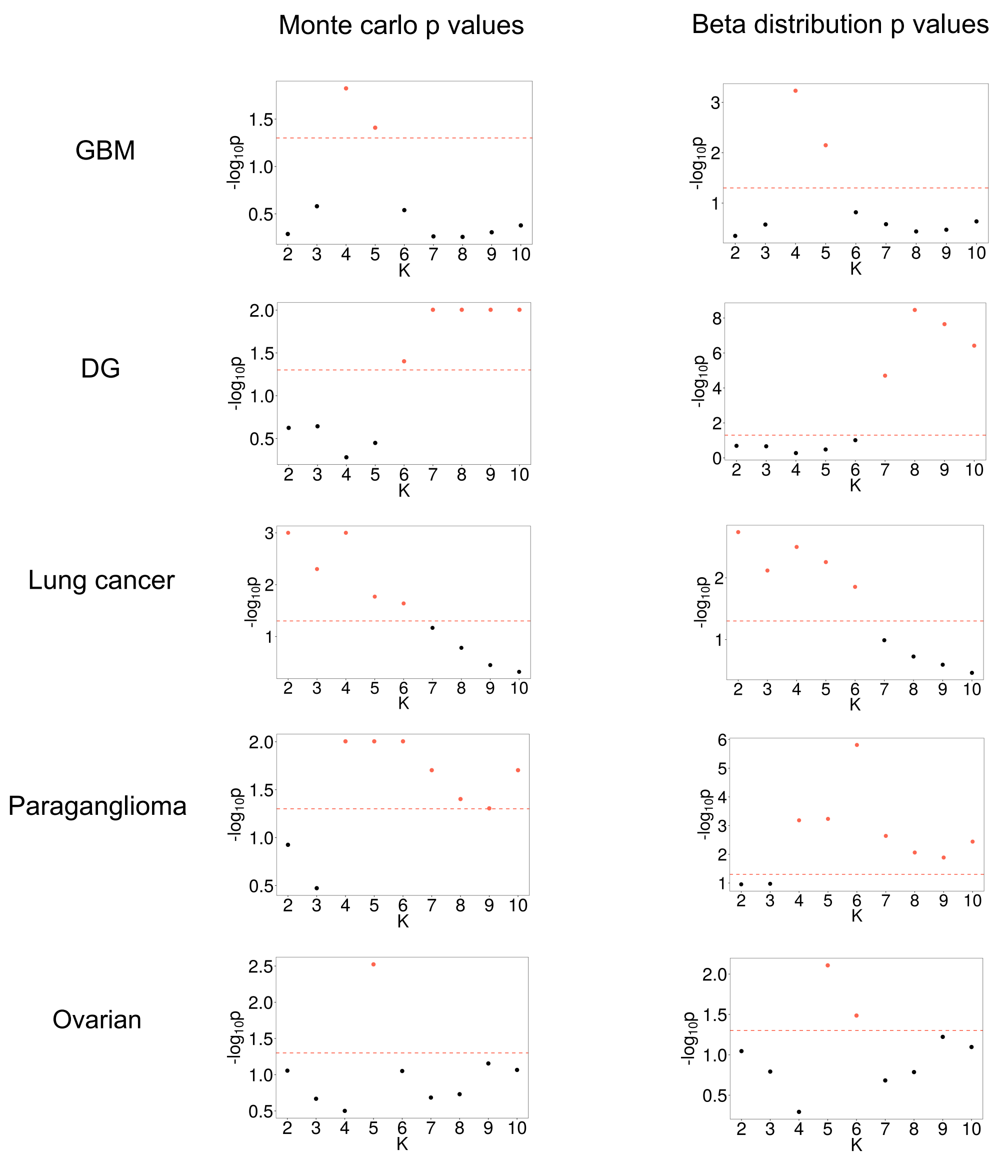

**Supplementary figure 5.** Comparing M3C Monte Carlo p values with p values estimated from a beta distribution. The beta distribution is generated using parameters from the initial $N$ Monte Carlo simulations M3C performs. Tail estimation allows the limits of a finite number of simulations to be overcome.

#### Supplementary figure 6

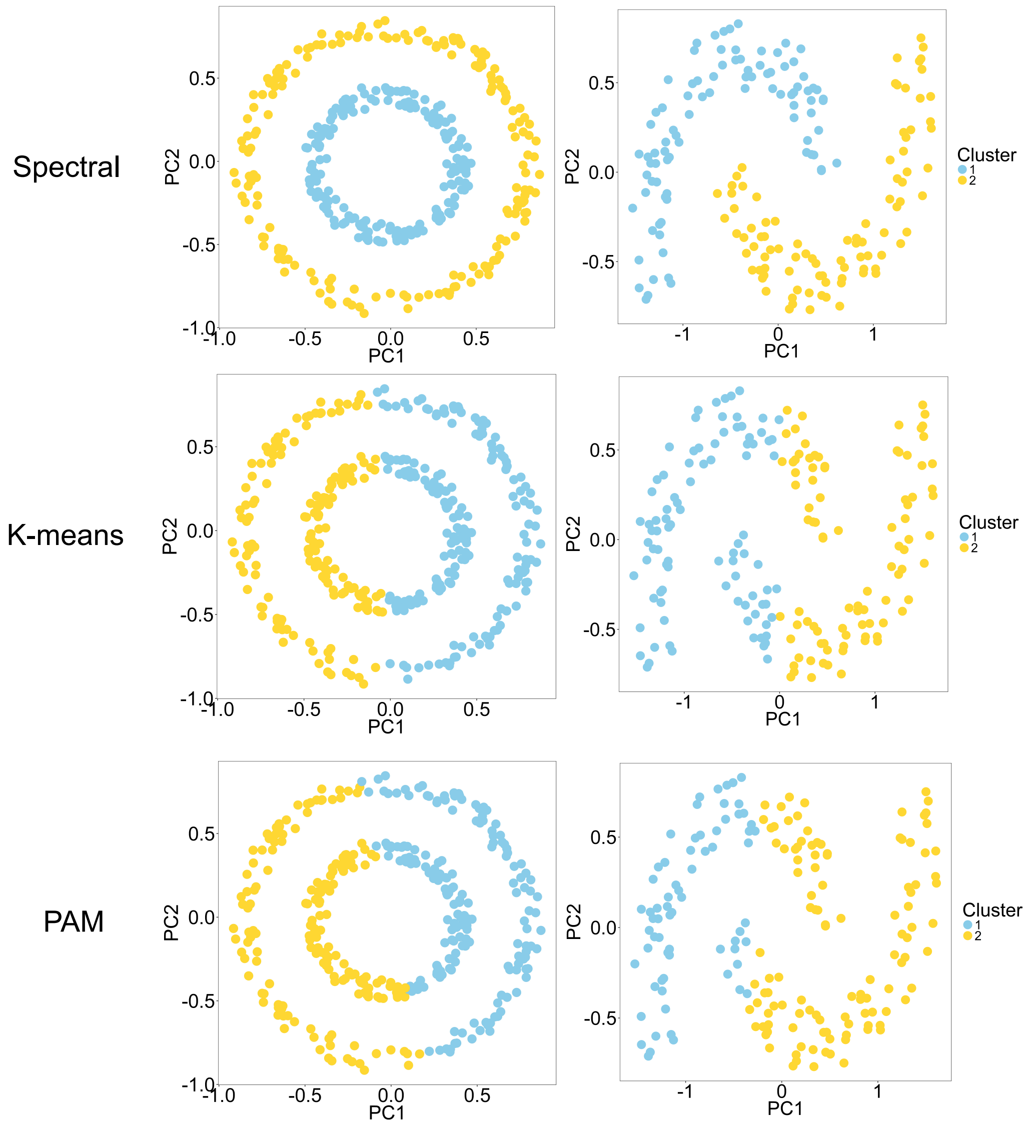

**Supplementary figure 6.** Comparing the M3C results using spectral clustering, PAM, or K-means on concentric circles or half-moon shapes.

**Supplementary figure 7**

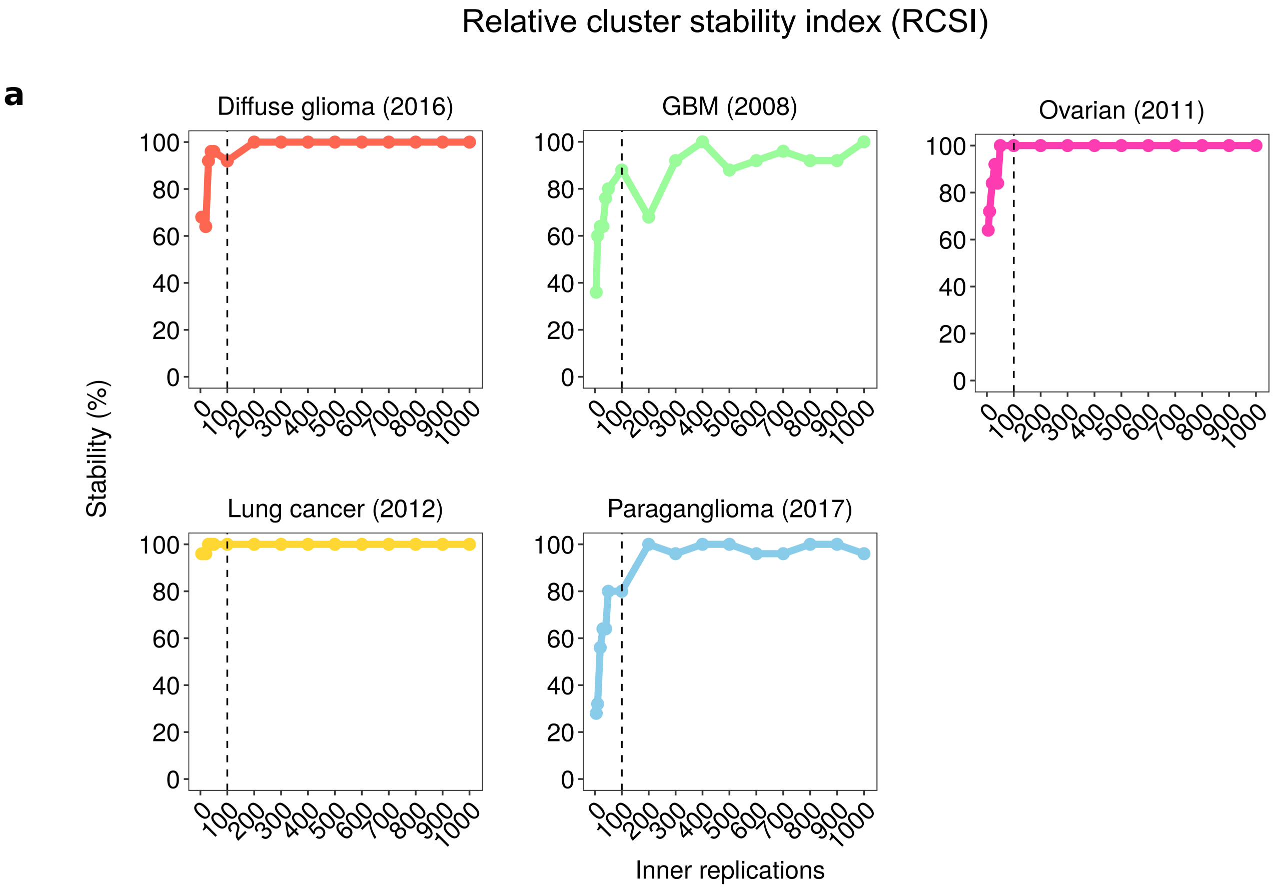


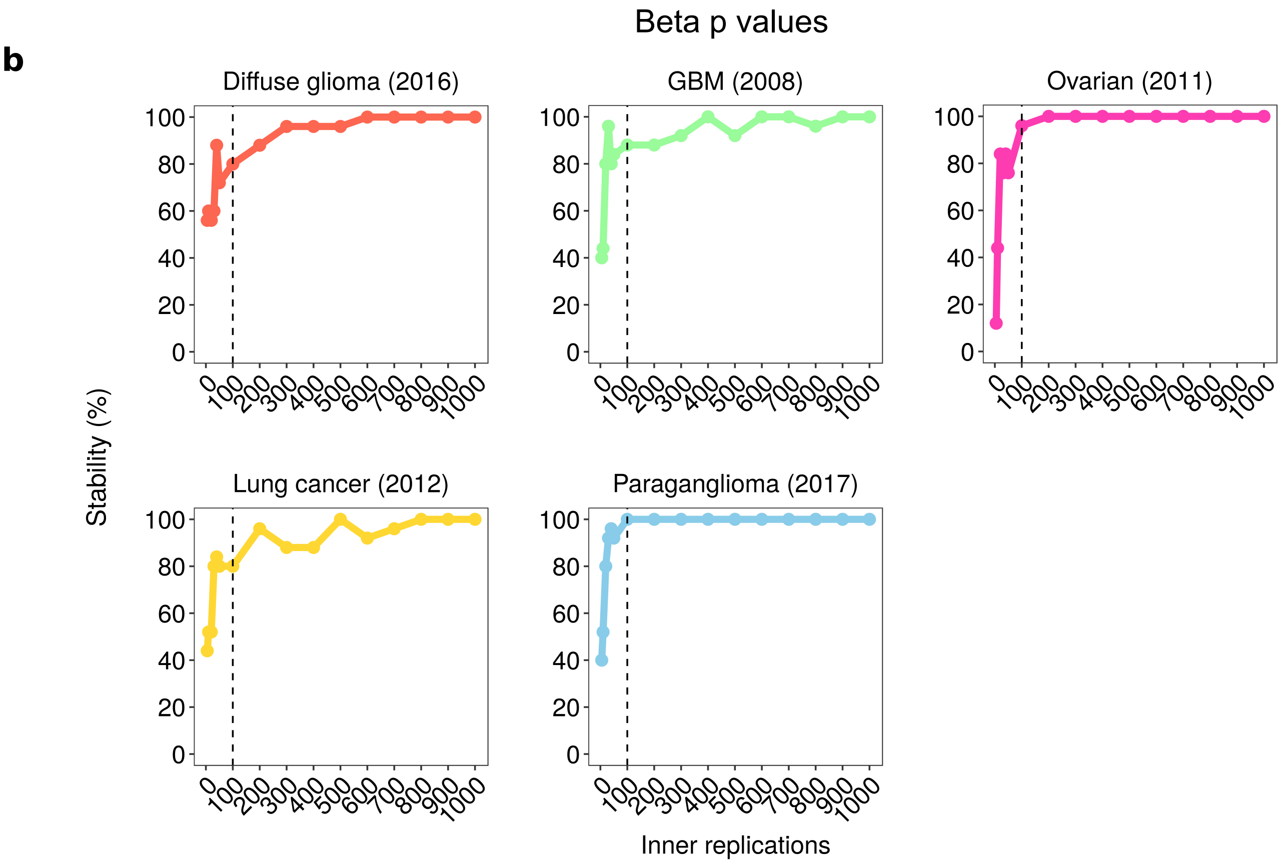

**Supplementary figure 7.** A sensitivity analysis of M3C parameters across 5 real datasets. (A) Results from the RCSI are shown. The inner resampling replications of M3C were varied, while the outer Monte Carlo simulations were left to default (value of 100). Stability is measured on the y axis as the percentage of times the same result was obtained across 25 iterations. (B) Results from the beta distribution after conducting the same analysis described in (A).

**Supplementary figure 8**

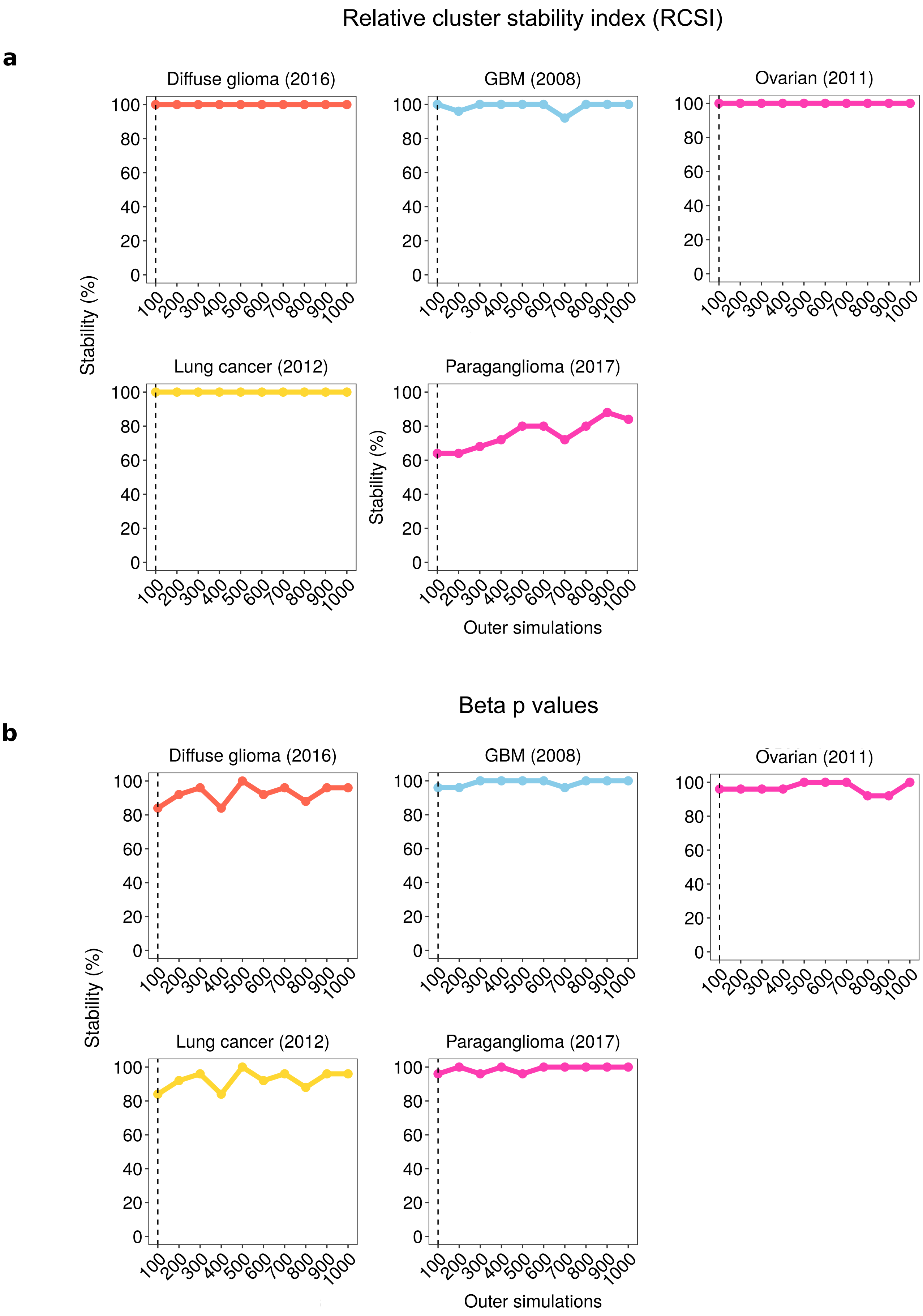

**Supplementary figure 8.** Same analysis described in supplementary figure 7, except the outer Monte Carlo simulations of M3C were varied, the inner resampling replications were left to default (value of 100).

**Supplementary Table I:** More detailed information about datasets selected for analysis. HC refers to hierarchical clustering.

| Publication | Year | Data type | Original Algorithm | Original score metric | Number of features | Data format | Samples | M3C K | Original K |
|---|---|---|---|---|---|---|---|---|---|
| Glioblastoma | 2008 | Microarray | Consensus clustering (HC) | Not stated | 1740 | log2(norm) | 206 | 4 | 4 |
| Ovarian Carcinoma | 2011 | Microarray | NMF Consensus clustering | Cophenetic | 800 | log2(norm) | 489 | 5 | 4 |
| Squamous cell lung cancers | 2012 | RNA-seq | Consensus clustering (HC) | Not stated | 2257 | log2(RPKM) | 178 | 2 | 4 |
| Human breast tumors | 2012 | miRNA-seq | NMF Consensus clustering | Cophenetic | 306 | log2(CPM) | 697 | 1 | 7 |
| Diffuse Glioma | 2016 | RNA-seq | Consensus clustering (HC) | Calinsky-Harabasz | 2275 | log2(CPM) | 667 | 8 | 4 |
| Lupus | 2016 | Microarray | HC | None | 48 | log2(norm) | 82 | 1 | 7 |
| Pheochromocytoma and Paraganglioma | 2017 | RNA-seq | Consensus clustering (HC) | Not stated | 3000 | log2(TPMs) | 173 | 6 | 4 |

**Supplementary note 1**

**M3C pseudo code**

INPUTS: $T$ = Data

PARAMS: $B$ = Monte Carlo simulations, $H$ = Inner resampling iterations, $K$ = maximum K, clustering algorithm = PAM (default)

#### REFERENCE

FOR $b = 1 \ldots B$:

  Calculate random data $Q^b$ (equations 1-3)

  Calculate Euclidean distance matrix

  FOR $h = 1 \ldots H$:

    Resample $Q^b$

    Create or update the indicator matrix I (equation 4)

    FOR $k = 1 \ldots K$:

      Cluster distance matrix using clustering algorithm

      Assign clustering to connectivity matrix $M$ (equation 5)

  Normalise $M$ (equation 6)

  Calculate CDF and reference PAC scores (equations 7-8)

#### REAL DATA

Calculate Euclidean distance matrix

FOR $h = 1 \ldots H$:

  Resample $T$

  Create or update the indicator matrix I (equation 4)

  FOR $k = 1 \ldots K$:

    Cluster distance matrix using inner clustering algorithm

    Assign clustering to connectivity matrix $M$ (equation 5)

Normalise $M$ for $k = 1 \ldots K$ (equation 6)

Calculate CDF then get real PAC score for each $k$ (equations 7-8)

#### FINAL STEPS

Calculate RCSI for $k = 1 \ldots K$ (equation 9)

Calculate p values for $k = 1 \ldots K$ (equation 10)
